# Pan-GWAS of *Streptococcus agalactiae* highlights lineage-specific genes associated with virulence and niche adaptation

**DOI:** 10.1101/574152

**Authors:** Andrea Gori, Odile Harrison, Ethwako Mlia, Yo Nishihara, Jacquline Chinkwita-Phiri, Macpherson Mallewa, Queen Dube, Todd D Swarthout, Angela H Nobbs, Martin Maiden, Neil French, Robert S Heyderman

**Affiliations:** NIHR Mucosal Pathogens Research Unit, Division of Infection and Immunity, University College London, 5 University Street, WC1E 6JF UK London, United Kingdom; Department of Zoology, University of Oxford, The Peter Medawar Building for Pathogen Research, OX1 3SY, Oxford, United Kingdom; Malawi-Liverpool-Wellcome Trust Clinical Research Programme, College of Medicine, University of Malawi, Blantyre, Malawi; University of Malawi, College of Medicine, Chichiri, Blantyre 3, Malawi; Queen Elizabeth Central Hospital, P.O. Box 95, Blantyre, Malawi; Liverpool School of Tropical Medicine, Clinical Sciences, Pembroke Place, L3 5QA, Liverpool, United Kingdom; Bristol Dental School, University of Bristol, Lower Maudlin Street, BS1 2LY, Bristol, United Kingdom; Department of Clinical Infection, Institute of Infection and Global Health, University of Liverpool, 8 West Derby Street, L69 7BE, Liverpool, United Kingdom

**Keywords:** *Streptococcus agalactiae*, pangenome, GWAS, population structure, bacterial phylogeny, virulence

## Abstract

*Streptococcus agalactiae* (Group B streptococcus, GBS) is a coloniser of the gastrointestinal and urogenital tracts, and an opportunistic pathogen of infants and adults. The worldwide population of GBS is characterised by Clonal Complexes (CCs) with different invasive potentials. CC17 for example, is a hypervirulent lineage commonly associated with neonatal sepsis and meningitis, while CC1 is less invasive in neonates and more commonly causes invasive disease in adults with co-morbidities. The genetic basis of GBS virulence and to what extent different CCs have adapted to different host environments remain uncertain. We have therefore applied a pan-genome wide association study approach to 1988 GBS strains isolated from different hosts and countries. Our analysis identified 279 CC-specific genes associated with virulence, disease, metabolism and regulation of cellular mechanisms that may explain the differential virulence potential of particular CCs. In CC17 and CC23 for example, we have identified genes encoding for pilus, quorum sensing proteins, and proteins for the uptake of ions and micronutrients which are absent in less invasive lineages. Moreover, in CC17, carriage and disease strains were distinguished by the allelic variants of 21 of these CC-specific genes. Together our data highlight the lineage-specific basis of GBS niche adaptation and virulence, and suggest that human-associated GBS CCs have largely evolved in animal hosts before crossing to the humans and then spreading clonally.

## INTRODUCTION

*Streptococcus agalactiae* (Group B *Streptococcus*, GBS) forms part of the normal gastrointestinal and urogenital microbiota, occasionally associated with causing life-threatening invasive disease in infants, pregnant women and adults with co-morbidities [Shabayek and Spellerberg, 2018]. Since the 1970s, GBS has been reported as one of the leading causes of neonatal mortality and morbidity in the US [Dermer, *et al*., 2004] but it is increasingly recognised that the burden is greatest in low-to-middle income countries. In sub-Saharan Africa, for example, where up to 30 percent of women carry GBS asymptomatically, the incidence of invasive GBS disease in neonates has been reported to be up to 2.1 per 1000 livebirths, with case fatality rates ranging from 13 to 46 percent [Dagnew, *et al*., 2012; Heyderman, *et al*., 2016; Nishihara *et al*., 2017].

In neonates, early-onset disease (EOD) in the first week of life typically presents as pneumonia or sepsis [Edmond et al., 2012, Nishihara *et al*., 2017]. Late-onset disease (LOD) develops from 7 days to 3 months after birth, and is frequently characterised by meningitis leading to chronic neurological damage, seizures, blindness and cognitive impairment in those that survive [Berardi et al., 2013; Nishihara *et al*., 2017]. The gastrointestinal tract is the reservoir for GBS and is the most likely source for maternal vaginal colonisation [Meyn *et al*., 2004]. This may lead to GBS transmission before or during birth, potentially leading to early onset disease in the infant [Nishihara *et al*., 2017]. The route for late onset colonisation and disease is less clear: while vertical transmission is still possible, environmental transmission and acquisition are considered more common [Rajagopal *et al*., 2009].

GBS capsular polysaccharide is a key virulence factor, mediating immune system evasion [Lemire *et al*., 2012], and is the basis for serotyping. Ten GBS capsular serotypes have been described [Slotved *et al*., 2007]. Serotypes Ia, Ib, II, III, and V account for 98% of human carriage serotypes isolated globally, although prevalence of each serotype varies by region [Russell *et al*., 2017]. Serotype III accounts for 25 to 30% of strains isolated in Europe and Africa but only 11% of strains isolated in Northern America or Asia. Serotypes VI, VII, VIII, and IX are frequently isolated in Southern, South-Eastern, and Eastern Asia but are relatively rare in other parts of the world [Russell *et al*., 2017]. Multi-locus sequence typing (MLST) has identified 6 major clonal complexes (CC) in humans: 1,10,17, 19, 23 and 26 [Da Cunha *et al*., 2014; Sørensen *et al*., 2014]. In recent years it has become apparent that some CCs have a greater potential to cause invasive disease, while others are largely associated with asymptomatic carriage. CCs 1, 23 and 19, for example, are the predominant colonisers of pregnant women, well adapted to vaginal mucosa with a limited invasive potential in neonates [Manning *et al*., 2008; Teatero *et al*., 2017]. In contrast, CC17 strains, mostly serotype III, are associated with neonatal sepsis and meningitis, and account for more than 80% of LOD [Lamy *et al*., 2006; Shabayek and Spellerberg., 2018]. Comparative phylogenetic analysis of human and bovine GBS strains suggested that CC17 emerged recently from a bovine ancestor (CC67) and is characterised by limited recombination [Bisharat *et al*., 2004]. However, this has been challenged [Shabayek and Spellerberg., 2018], and the relationship between isolates from these different hosts remains uncertain.

Colonisation and persistence of GBS in different host niches is dependent upon the ability of GBS to adhere to the mucosal epithelium [Shabayek and Spellerberg., 2018; Nobbs *et al*., 2009; Rosini and Margarit, 2015], utilising numerous bacterial adhesins including fibrinogen binding protein (Fbs), the group B streptococcal C5a peptidase (ScpB) and the GBS immunogenic bacterial adhesin (BibA) [Landwehr-Kenzel and Henneke, 2014; Cheng *et al*., 2002; Santi *et al*., 2006]. Biofilm formation is essential to promoting colonisation, which is also enhanced by bacterial capsule and type IIa pili [Konto-Ghiorghi *et al*., 2009; Xia *et al*., 2015]. Biofilm formation also plays a central role in the phenotype switch from commensal to pathogen [Patras *et al*., 2018]. Recently, deletion of the gene for Biofilm regulatory protein A (BrpA) was shown to impair both the biofilm formation and the ability of the bacterium to colonise and invade the murine host [Patras *et al*., 2018]. The expression of these virulence factors vary by CC, with the Fbs proteins carried by the hypervirulent lineage CC17, for instance, characterised by specific deletions and frameshift mutations that alter the sequence or expression rate [Buscetta *et al*., 2014]. *S. agalactiae* is able to survive both the acidic vaginal environment and within the blood [Santi *et al*., 2009]. Transcription analyses have suggested that this transition is largely mediated by two component system CovRS [Patras *et al*., 2013; Almeida *et al*., 2015]. Recently, specific gene substitutions in TCS CovRS have been identified in disease-adapted CC17 GBS clones [Almeida *et al*., 2017]. How widespread these genetic adaptations are amongst CC17 and whether different adaptations confer enhanced colonisation and disease potential in other CCs is uncertain.

Here, we report a pan-genome wide association study of genome sequence data from 1988 GBS carriage or invasive disease isolates from different hosts and countries. This revealed that GBS CCs possessed distinct collections of genes conferring increased potential for persistence including genes associated with carbohydrate metabolism, nutrient acquisition and quorum-sensing. Within CC17, allelic variants of these crucial genes distinguish carriage from invasive strains. The differences in the GBS CCs analysed are not geographically restricted, but may have emerged from an original ancestral GBS strain in animal hosts before crossing to humans.

## METHODS

### Bacterial strains, genomes and origin

Publicly available genome sequences from 1574 human isolates from Kenya, USA, Canada and the Netherlands, together with 111 genomes from animal isolates were analysed (Seale *et al*., 2015; Flores *et al*., 2015; Teatero *et al*., 2014; Jamrozy *et al*., 2018; Table 1). The genome assemblies were not available for the isolates from Kenya and the Netherlands. In those cases, short read sequence data were retrieved from the European Nucleotide Archive (ENA, https://www.ebi.ac.uk/ena). Raw DNA reads were trimmed of low-quality ends and cleaned of adapters using Trimmomatic software (ver. 0.32; Bolger *et al*., 2014) and a sample of 1400000 reads for each paired-end library (e.g. 700000 reads x 2) was used for de-novo assembly. De-novo assembly was performed with SPAdes software (ver 3.8.0, Bankevich *et al*., 2012), using k-mer values of 21, 33, 55 and 77, automatic coverage cutoff, and removal of contigs 200 bp-long or shorter. De-novo assemblies were checked for plausible length (between 1900000 and 2200000 bp), annotated using Prokka (ver. 1.12; Seemann, 2014) and checked for low-level contamination using Kraken software (ver. 0.10.5; Wood and Salzberg, 2014). In cases for which more than 5% of the contigs belonged to a species different from *Streptococcus agalactiae*, the genome sequence was flagged as contaminated and not included in any further analysis. Resulting assemblies were deposited in the pubmlst.org/sagalactiae database which runs the BIGSdb genomics platform (Jolley and Maiden, 2006).

**Table 1 –.** Characteristics of GBS isolates. Animal isolates are reported to be isolated from cattle (n=83), fish (n=24) and frogs (n=3). * Isolated from Queen Elizabeth Central Hospital, Blantyre; ** Jamrozy, *et al*., 2018; *** genomes retrieved from pubmlst.org. Full metadata are reported in Supplementary table S1.

|  | Country |  | Count | # Invasive | # Missing data |
| --- | --- | --- | --- | --- | --- |
| **Human isolates** | Malawi | This work* | 303 | 131 | 6 |
|  | Kenya | Seale et al., 2015 | 1034 | 71 | 0 |
|  | USA | Flores et al., 2015 | 99 | 99 | 0 |
|  | Canada | Teatero et al., 2014 | 141 | 141 | 0 |
|  | The Netherlands | PRJEB14124** | 300 | unknown | 300 |
| **Animal isolates** | Italy | *** | 3 |  |  |
|  | Kenya | *** | 2 |  |  |
|  | Germany | *** | 1 |  |  |
|  | Brazil | *** | 1 |  |  |
|  | Unknown | *** | 104 |  |  |

In addition, 303 carriage and invasive disease strains isolated in Malawi between 2004 and 2016 in the context of carriage and invasive disease surveillance were sequenced. For these, DNA was extracted from an overnight culture using DNAeasy blood & tissue kit (Qiagen^®^) following manufacturer’s guidelines for bacterial DNA, and sequenced using HiSeq4000 (paired-end library 2×150) platform at Oxford Genomics Centre UK. Sequences were then assembled as described above.

### MLST and Serotype definition

Serotypes were determined via DNA sequence similarity, as described previously [Seale, *et al*. 2016]. BLASTn was used to align the DNA fragments typical of each serotype to the DNA assemblies of the isolates with the following parameters: evalue 1e-10, minimum 95 percent identity, minimum 90% query coverage, and the results for each BLASTn alignment was parsed with ad-hoc perl scripts. Accession numbers for the sequences used were AB028896.2 (from 6982 to 11695, serotype Ia); AB050723.1 (from 2264 to 6880, serotype Ib); EF990365.1 (from 1915 to 8221, serotype II); AF163833.1 (from 6592 to 11193, serotype III); AF355776.1 (from 6417 to 11656, serotype IV); AF349539.1 (from 6400 to 12547, serotype V); AF337958.1 (from 6437 to 10913, serotype VI); AY376403.1 (from 3403 to 8666, serotype VII); AY375363.1 (from 2971 to 7340, serotype VIII). Only one fragment matched each genome under these parameters, and it defined each isolate’s serotype. If none of the serotype defining DNA fragments matched under the described parameters, the isolate was defined as Non-Typeable (NT).

Multi-locus sequence types (MLST) STs were derived from the allelic profiles of the 7 housekeeping genes (*adhP, pheS, atr, glnA, sdhA, glcK tkt*). This grouped strains into 91 unique STs. Strains which did not show a full set of housekeeping gene alleles or were not assigned to any previously described ST (n=68) were double-checked for sequence contamination and assigned to a non-sequence typeable (NST) group.

### Phylogeny inference

BURST [Enright *et al*., 2002] was used to evaluate the relatedness between different STs, and to define CCs. Five random subsets, each containing 1000/1988 isolates, were analysed using eBURST on PubMLST [Jolley and Maiden, 2006]. This grouped STs sharing at least five out of seven MLST loci, and identified the central ST (i.e. the ST with the highest number of single or double locus variants), and was used to define CCs. Each of the five subsets showed the same six CCs (CC1, CC6, CC10, CC19, CC17 and CC23) plus a series of singletons (STs not belonging to any CC). CCs were defined as the set of STs associated with a particular CC in at least one run of eBURST.

Core-genome phylogeny of GBS datasets was inferred using the software Parsnp (from Harvest package, ver. 1.1.2; Treangen *et al*., 2014), which performs a core genome SNP typing and uses Fastree2 [Price *et al*., 2010] to reconstruct whole-genome maximum-likelihood phylogeny, under a generalised time-reversible model. Each tree shown was rooted at mid-point. Parsnp requires a reference to calculate the core SNPs shared by all isolates: complete finished reference genomes from 5 different strains were used separately (accession numbers: NC_021485 – strain 09mas018883 – CC1; NC_007432 – strain A909 – CC6; HG939456 – strain COH1 – CC17; NC_018646 - strain GD201008 – CC6; NC_004368 – strain NEM316 – CC23). Visualisation of the phylogenetic analysis was performed via iTol (Letunic and Bork, 2016)

### Pangenome construction and genome wide association analysis

A pangenome was generated from the combined African (isolates from Malawi and Kenya, Seale *et al*., 2016), Canadian [Teatero *et al*., 2014], American [Flores *et al*., 2015], Dutch and animal-derived strains using Roary, (ver. 3.8.0; Page *et al*., 2015). Parameters for each run were: 95% of minimum blastp identity; MLC inflation value 1.5; with 99% as the percentage cutoff in which a gene must be present to be considered as core.

In the last decade, several pipelines have been developed for bacterial genome wide association studies (GWAS), such as PLINK, PhyC, ROADTRIPS and SEER [Chen and Shapiro, 2015; Chang *et al*., 2015; Thornton *et al*., 2010; Lees *et al*., 2016]. Scoary [Brynildsrud *et al*., 2016] was designed to highlight genes in the accessory pangenome of a bacterial dataset associated with a particular bacterial phenotype: it can deal with either binary/discrete phenotypes (+/− e.g. bacterial colony colour) or continuous phenotypes (e.g. antimicrobial resistance). In this analysis, Scoary (ver. 1.6.16) was used to establish which genes were typical of each CC via a Pangenome-Wide Association Study (pan-GWAS). The CC of each isolate was depicted as a discrete phenotype, e.g. belonging to CC17 or not, and defined as “positive” or “negative” respectively with the Scoary algorithm evaluating which gene feature is statistically associated with a particular CC [Brynildsrud *et al*., 2016]. The cut-off for a significant association was a *p*-value lower than 1e-10 and a sensitivity and specificity greater than 90 percent. Genes associated with CC1, CC10, CC19, CC17 or CC23 were plotted on the circular representation of the chromosome of 5 GBS isolates belonging to each CC (strains ST-1; NCTC8187; 2603V/R; SGM4; 874391; NGBS572). The plot was obtained with BRIG (ver. 0.8; Alikhan *et al*., 2011). Gene syntheny was then evaluated for those genes found to be associated with each CC. To do this, three genomes belonging to each CC were selected, aligned using ProgressiveMauve and the genes identified from the pan-GWAS analysis plotted [Darling *et al*., 2010]. Mauve (ver. 2.3.1; Darling *et al*., 2004) was used to produce a graphical representation of the alignment and gene syntheny was qualitatively evaluated.

Sequence diversity of genes identified from the pan-GWAS analysis was investigated by selecting one representative nucleotide gene sequence associated with each CC (sequences reported in supplementary information file 1) and aligning this against each genome included in the analysis using BLASTn version 2.3. The bitscore value of each gene, aligned against each isolate, was used to produce the heatmap shown in Supplementary figure S2, using the R package pheatmap (ver. 1.0.10; https://CRAN.R-project.org/package=pheatmap). Bitscores were normalised against (i.e. divided by) the highest scoring isolate for each gene: the normalised bitscore was 0>x>=1 where 1 corresponds to the highest identified bitscore, 0 corresponds to the absence of the gene, and values in between highlight a different level of gene similarity. For the identification of alleles that distinguish strains isolated from disease from carriage, we calculated the allelic profiles of the genes identified by the pan-GWAS pipeline in the 547 CC 17 strains (for which the source of isolation was non-animal and known). For each gene we selected the alleles present in at least 10 strains, and calculated the proportion of strains isolated from invasive source and carriage. Significance for alleles unevenly distributed between carriage and disease was calculated with Fisher test.

### Ethical approval for Malawi GBS Collection

Collection of carriage isolates was approved by College of Medicine Research Ethics Committee (COMREC), University of Malawi (P.05/14/1574) and the Liverpool School of Tropical Medicine Research Ethics Committee (14.036). Invasive disease surveillance in was approved by COMREC (P.11/09/835 and P.08/14/1614).

## RESULTS

### GBS whole genome sequencing dataset

A total of 358 Malawian GBS genome sequences were initially available. Of these, 55 samples did not pass the quality checks, therefore the final Malawian dataset was composed of 303 GBS strains (Table 1), including 131 isolated from invasive disease in children and 166 isolated from healthy mothers, in which draft genome assemblies had an average N50 of 163462 (range 12593 – 717849), average contigs number of 70 (range 20 – 377) and average longest contig of 30148 (range 44691 – 1019176).

Five further datasets were included (1674 clinical isolates) composed of 1034 Kenyan strains Kenya, 99 American, 141 Canadian, and 300 Dutch strains randomly selected from 1512 isolates from the Netherlands (Table 1). A total of 111 animal isolates sampled in several different countries was also included (Table 1). Where information was available, 446 (22.4% of the 1988 total) strains were associated with invasive disease (bacteraemia or meningitis) and 1125 (56.6%) from healthy carriers. Meta-data consisting of country of origin, year of isolation, capsular serotype, MLST-ST and accession number are reported in Table S1. The genome sequences from 1998 isolates were used for the analysis.

### Clonal-complex assignment and core genome phylogeny

Six CCs were identified: CC1, CC6, CC10, CC19, CC17 and CC23 according to the groups defined using the BURST algorithm [Enright *et al*., 2002] and core genome phylogeny (Table S1 and S2). Several STs were found in just one country (e.g. ST 866 found exclusively in Malawi or ST 196 in Kenya); however, these STs were always represented by less than 20 isolates. With the exception of the USA, where isolates were intentionally selected to represent only CC1 [Teatero *et al*., 2014], and the rare CC6 represented by 22 isolates, CCs were distributed across all of the countries analysed.

While each serotype in the clinically derived WGS was predominantly associated with only one or two CCs, the animal isolates were more variable (Figure 1). SNPs identified in the part of the genome shared by all isolates (~26000 polymorphisms) were used to infer the ML phylogenetic trees (Figure 2). CCs clustered in distinct branches of the tree; in particular, CC17 and CC23 produced two clusters. Although the majority of animal derived WGS data clustered within a separate branch, 59/111 isolates were located in clusters that were associated with human derived samples and CCs. For example, three animal isolates (LMG15085, LMG15094 and CI7628) clustered with the clinical isolates in the CC17. This pattern raises the possibility that the human-associated CCs analysed here arose in animals and then underwent zoonotic transfer and clonal expansion after infection of the human host.

**Figure 1 –.**
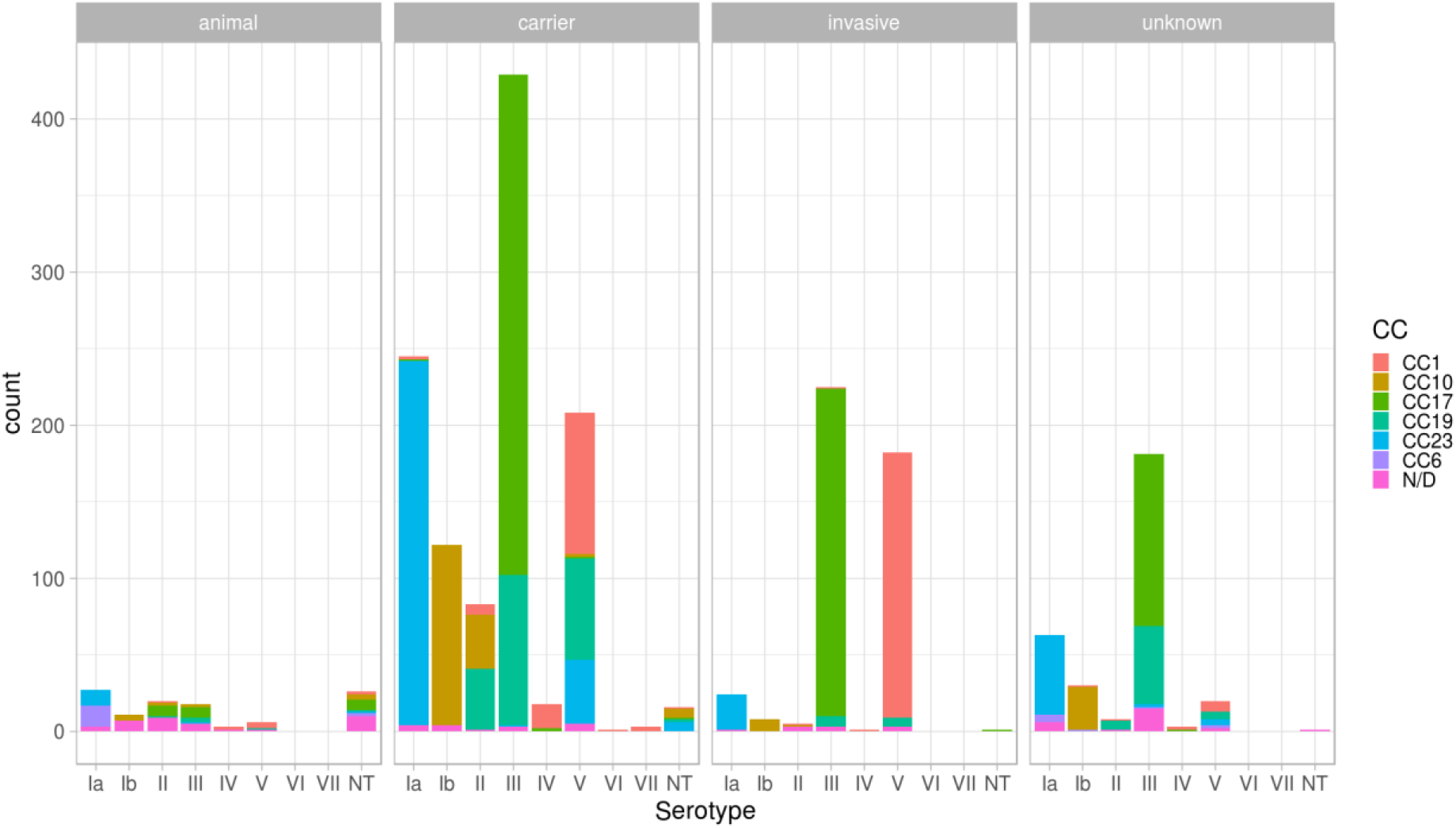
Isolates used in this study, stratified per serotype, CC and source. Clinical isolates are grouped as “invasive” (including strains isolated from children and adults affected by any GBS invasive disease), “carrier” (including healthy carrying mothers), and “unknown” where metadata were not available.

**Figure 2 –.**
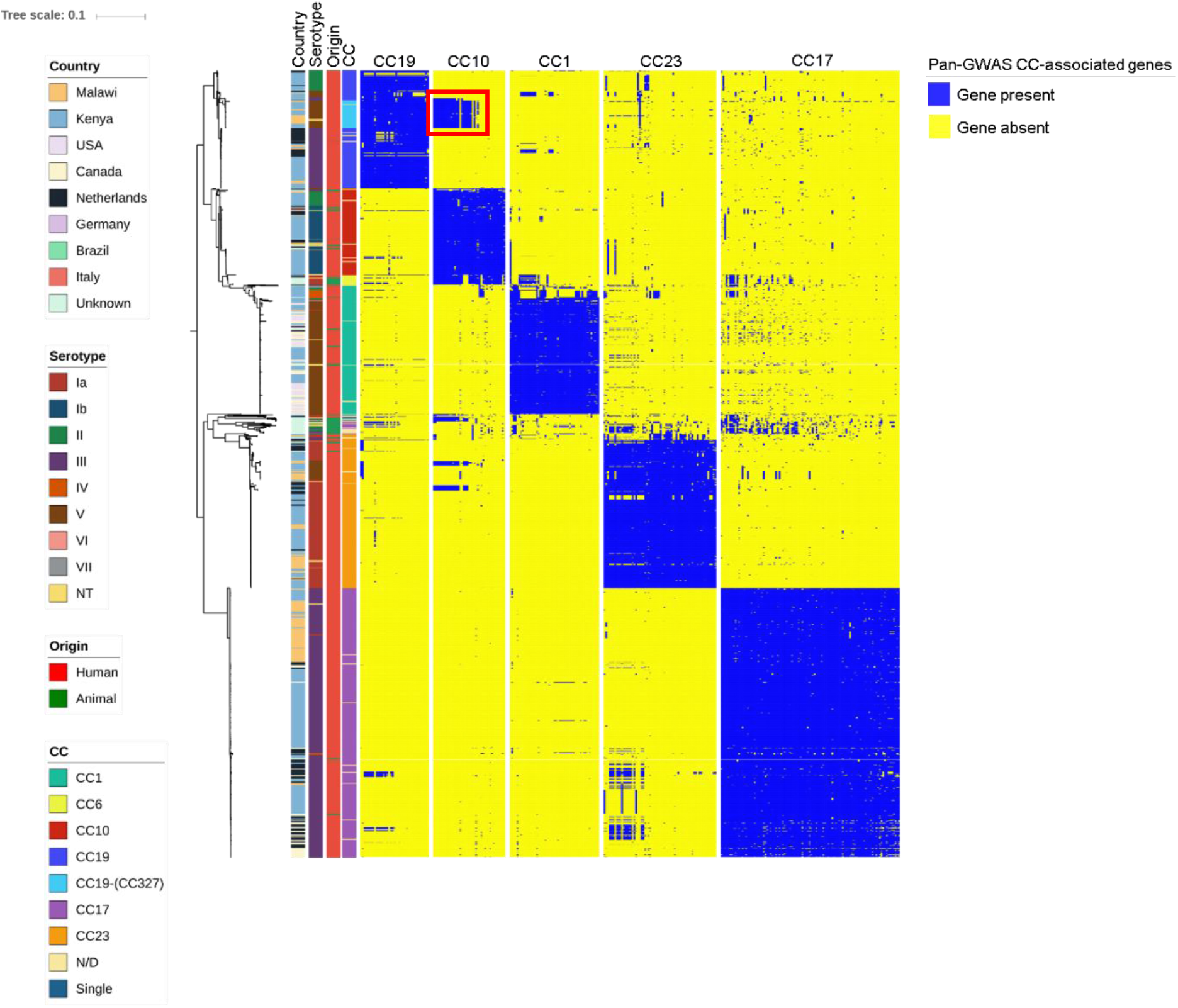
Core-genome based population structure of GBS. The phylogenetic tree is annotated with 4 coloured strips representing the clonal complex, the country of isolation, the origin and the serotype of each strain. The three binary heatmaps, represent the presence (blue) or absence (yellow) of the genes identified by the pan-GWAS pipeline. Tree is rooted at midpoint. The reference strain used in this analysis was COH1 - reference HG939456. The red square in the “CC10” heatmap highlights the cluster of CC10-associated genes found in CC19 clones. Trees build with different reference strains are shown in figure S1, and show analogous topology.

### Pangenome and pan-GWAS

Scoary has previously been used for a similar pan-GWAS analysis of 3 CC17 strains [Almeida *et al*., 2017]. In this study, we applied it to a pangenome built on a dataset of 1988 strains, representing 6 different clonal complexes (Figure 1): according to the roary nomenclature [Page *et al*., 2015], 1374 genes were included in the core genome (i.e. present in more than 95% of the strains, “core” and “soft-core” genes), and 12457 genes in the accessory genome (i.e. less than 95% of the strains, “shell” and “cloud” genes). We observed that saturation of the pangenome was achieved. A total of 51, 41, 39, 102 and 64 genes associated with CC1, CC10, CC19, CC17 and CC23 respectively (Table 2; Table S3) were identified, with a specificity and sensitivity in defining the CC given the annotated CDS and vice-versa greater than 90% (p<0.05). The pipeline was not applied to CC6, which was represented by only 22 genomes in our dataset. BLASTn was used to confirm whether gene sequences associated with each CC in the pan-GWAS were completely absent in different CCs, or had accumulated sufficient mutations to fail recognition by automated annotation (i.e. PROKKA). We identified 57 such genes in CC17 out of the 102 identified by the Scoary pipeline, 22 genes in CC23, 4 genes in CC1, 9 genes in CC10 and 5 in CC19 (Figure S2; Table S3). This suggests that the genes characterising a particular CC may have been rendered non-functional (i.e. as pseudogenes) in other CCs (Table S3 highlights which CC-associated genes are completely absent, and which genes are characterised by mutations - SNPs or In-dels - that alter the protein sequence with point mutations or truncation).

**Table 2 –.** Pathways and functional categories identified by KEGG annotation in the five groups of CC-associated genes. For each clonal complex the functional category pathway is shown on the right-end side of the table. For each functional category, the metabolic pathway affected and its Kegg reference number are reported.

| **CC1** | **Kegg #** | **Pathway** |
| --- | --- | --- |
| **Metabolism (09100)** | 01130 | Biosynthesis of antibiotics |
|  | 00052 | Galactose metabolism |
|  | 00999 | Biosynthesis of secondary metabolites - unclassified |
| **Environmental Information Processing (09130)** | 02060 | Phosphotransferase system (PTS) |
| **CC10** |  |  |
| **Metabolism (09100)** | 01100 | Metabolic pathways |
|  | 01110 | Biosynthesis of secondary metabolites |
|  | 01120 | Microbial metabolism in diverse environments |
|  | 01130 | Biosynthesis of antibiotics |
|  | 00010 | Glycolysis / Gluconeogenesis |
|  | 00040 | Pentose and glucuronate interconversions |
|  | 00051 | Fructose and mannose metabolism |
|  | 00052 | Galactose metabolism |
|  | 00561 | Glycerolipid metabolism |
|  | 00600 | Sphingolipid metabolism |
|  | 00603 | Glycosphingolipid biosynthesis - globo and isoglobo series |
| **Environmental Information Processing (09130)** | 02060 | Phosphotransferase system (PTS) |
| **CC19** |  |  |
| **Metabolism (09100)** | 01100 | Metabolic pathways |
|  | 00270 | Cysteine and methionine metabolism |
|  | 00760 | Nicotinate and nicotinamide metabolism |
| **Environmental Information Processing (09130)** | 03070 | Bacterial secretion system |
| **CC17** |  |  |
| **Metabolism (09100)** | 00010 | Glycolysis / Gluconeogenesis |
|  | 00020 | Citrate cycle (TCA cycle) |
|  | 00052 | Galactose metabolism |
|  | 00500 | Starch and sucrose metabolism |
|  | 00520 | Amino sugar and nucleotide sugar metabolism |
|  | 00620 | Pyruvate metabolism |
|  | 00630 | Glyoxylate and dicarboxylate metabolism |
|  | 00640 | Propanoate metabolism |
|  | 00680 | Methane metabolism |
|  | 00910 | Nitrogen metabolism |
|  | 00561 | Glycerolipid metabolism |
|  | 00230 | Purine metabolism |
|  | 00240 | Pyrimidine metabolism |
|  | 00250 | Alanine, aspartate and glutamate metabolism |
|  | 00260 | Glycine, serine and threonine metabolism |
|  | 00280 | Valine, leucine and isoleucine degradation |
|  | 00220 | Arginine biosynthesis |
|  | 01007 | Amino acid related enzymes |
|  | 00430 | Taurine and hypotaurine metabolism |
|  | 01003 | Glycosyltransferases |
|  | 01005 | Lipopolysaccharide biosynthesis proteins |
|  | 01011 | Peptidoglycan biosynthesis and degradation proteins |
|  | 00760 | Nicotinate and nicotinamide metabolism |
|  | 00770 | Pantothenate and CoA biosynthesis |
|  | 01001 | Protein kinases |
|  | 01002 | Peptidases |
|  | 03021 | Transcription machinery |
|  | 03016 | Transfer RNA biogenesis |
| **CC17 (continue)** |  |  |
|  | 00970 | Aminoacyl-tRNA biosynthesis |
|  | 03110 | Chaperones and folding catalysts |
|  | 03060 | Protein export |
|  | 03420 | Nucleotide excision repair |
|  | 03400 | DNA repair and recombination proteins |
| **Environmental Information Processing (09130)** | 02000 | Transporters |
|  | 02010 | ABC transporters |
|  | 02060 | Phosphotransferase system (PTS) |
|  | 03070 | Bacterial secretion system |
|  | 02020 | Two-component system |
|  | 02044 | Secretion system |
|  | 02022 | Two-component system |
| **Cellular Processes (09140)** | 04147 | Exosome |
|  | 02048 | Prokaryotic Defense System |
|  | 02024 | Quorum sensing |
|  | 02026 | Biofilm formation - Escherichia coli |
| **Unclassified (09190)** | 99982 | Energy metabolism |
|  | 99984 | Nucleotide metabolism |
|  | 99999 | Others |
|  | 99977 | Transport |
| **CC23** |  |  |
| **Metabolism (09100)** | 00630 | Glyoxylate and dicarboxylate metabolism |
|  | 00061 | Fatty acid biosynthesis |
|  | 01040 | Biosynthesis of unsaturated fatty acids |
|  | 01004 | Lipid biosynthesis proteins |
|  | 00260 | Glycine, serine and threonine metabolism |
|  | 00550 | Peptidoglycan biosynthesis |
|  | 01011 | Peptidoglycan biosynthesis and degradation proteins |
|  | 00780 | Biotin metabolism |
|  | 00670 | One carbon pool by folate |
|  | 01008 | Polyketide biosynthesis proteins |
|  | 01053 | Biosynthesis of siderophore group nonribosomal peptides |
|  | 00333 | Prodigiosin biosyntheses |
|  | 01002 | Peptidases |
| **Cellular Processes (09140)** | 02000 | Transporters |
|  | 02010 | ABC transporters |
|  | 02020 | Two-component system |
|  | 02042 | Bacterial toxins |
| **Human Disease (09100)** | 02048 | Prokaryotic Defense System |
|  | 02024 | Quorum sensing |
|  | 01502 | Vancomycin resistance |
|  | 01504 | Antimicrobial resistance genes |
| **Unclassified (09190)** | 99988 | Biosynthesis and biodegradation of secondary metabolites |
| **Genetic Information Processing (09120)** | 03000 | Transcription factors |

Gene location identified from the pan-GWAS analyses in CC1, CC10, CC17 and CC23 was evenly spread across the chromosome, and not clustered in a particular area consistent with the gene associations observed not resulting from a chromosomally integrated plasmid or transposon pathogenicity island acquired through horizontal gene transfer (Figure 3). One exception was CC19, where the majority of the 39 genes were clustered in 200 kbp region of the chromosome. Gene syntheny was conserved across different isolates (Gene syntheny in CC17 isolates is shown in figure S3).

**Figure 3 –.**
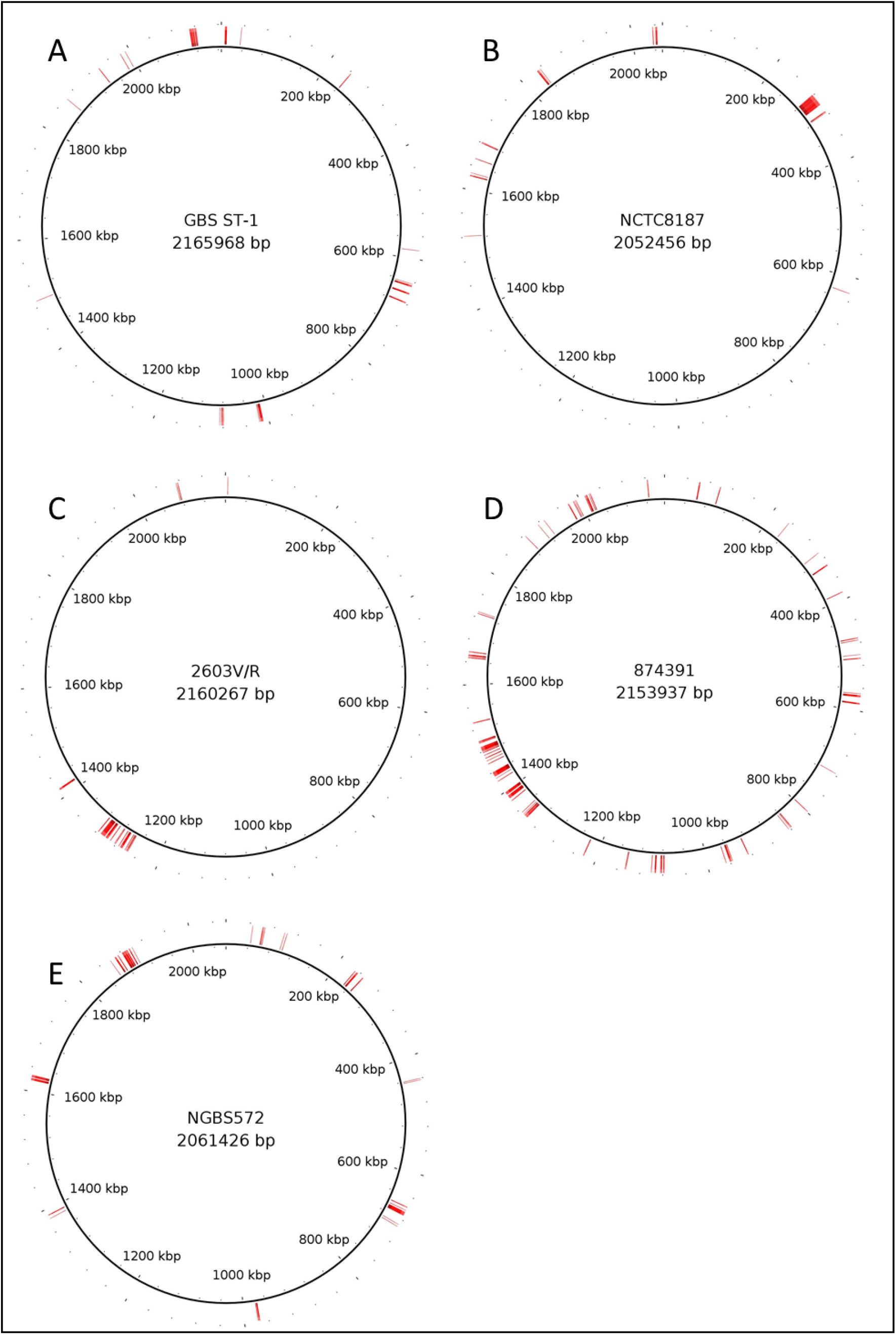
Location of genes identified by the pan-GWAS pipeline on a strain belonging to CC1 (A), CC10 (B), CC19 (C), CC17 (D) and CC23 (E). Gene location on each chromosome is represented by a red mark.

The majority of the pan-GWAS identified genes were associated with only one CC, but a particular cluster of genes associated with CC10 (including the *gatKTEM* system for galactose metabolism) was also present in a set of isolates belonging to CC19 (Figure 2). These isolates were all from Africa (Malawi and Kenya) and were ST-327 and ST-328.

### Functional pathways affected by CC-specific genes

A total of 279 genes were found to be CC-specific (Table S3). Genes characteristic of CC17 and CC23 were classified into five functional categories (Table 2): metabolism, environmental information processing, cellular processes, human disease, genetic information processing. In both CCs, the most represented functional families were those including metabolic genes and environmental information processes.

Differences in metabolic pathways between CC17 and CC23 included carbohydrate, amino acid, nitrogen compound and fatty acid metabolism. Siderophores for the uptake and transport of micronutrients (i.e. iron or nickel), and essential for successful colonisation of the human host in several bacterial pathogens [Bray *et al*., 2009; Janulczyk, *et al*., 2003; Kehl-Fie *et al*., 2013], also exhibited significant variation, for instance with genes for nickel uptake (*nikE* and *nikD*) and iron transport (*feuC*) truncated or characterised by SNPs in non-CC17 strains (Table S3).

CC17 and CC23 also showed differences in the genes affecting the environmental information processing functional pathways characterised by the presence of phosphotransferase (PTS) systems and two component systems (TCS), used for signal transduction and sensing of environmental stimuli. Moreover, in the same functional category, differences were present in secretion systems, transporters, quorum sensing and bacterial toxins. These pathways are used by GBS not only in colonisation of the host, but also to gain competitive advantage with other microorganisms occupying a particular ecological niche [Paterson *et al*., 2006].

Genes for prokaryotic defence systems, such as the CRISPR-Cas9 system, were also found, as well as proteins involved in genetic information processing such as transcription factors and regulators that may affect the expression of multiple genes [Lier *et al*., 2015]. Finally, antibiotic resistance also appears amongst the lineage specific characteristics; in particular, CC23 is the only CC showing typical genes involved in vancomycin resistance. CC17 also showed the presence of genes belonging to the KEGG group for “Nucleotide excision repair” and “DNA repair/recombination protein” (KO numbers 03420/03400, Table 2) which could indicate a variation in mutagenesis rate, thus capacity to respond to changes in environmental conditions and presence of stresses.

In contrast, the genes defining CC1, 10 and 19 were confined to metabolism, environmental information processing and genetic information processing. Genes involved with regulation and environmental sensing (PTS systems), as well as secretion systems were identified in this group of CCs. In particular, a gene encoding for the VirD4 type IV secretion system protein was associated with CC19. CC10 was characterised by an array of genes involved in carbohydrate metabolism and uptake, such as the ABC transport system for multiple sugar transport.

The majority of genes characteristic for CC1 were of unknown function, with the exception of genes involved with genetic regulation and a complete toxin/antitoxin system *phd/doc* [Chan *et al*., 2014]. These systems are often described as a tool for stabilising extrachromosomal DNA (i.e. plasmids), but they are often found integrated chromosomally in both Gram positive and Gram negative bacterial species, and their function when in this setting is unclear [Van Melderen, 2010].

In relation to the CC17-associated genes, we also checked for allelic variants specific to strains isolated from invasive disease or carriage. Figure S4 shows the proportion of CC17 invasive or carriage strains, and the frequency of each allelic variant. We identified 21 genes with alleles that statistically differentiated strains isolated from carriage and invasive disease (Fisher test, p<0.05, Table3. The DNA sequence of the allelic variant differed by a single polymorphism in all cases. In 15/21 cases this nucleotide change was translated into an aminoacid change (missense mutation), while in only a single case the mutation was nonsense, resulting in the truncated protein. This was the case of *gcc1730*, encoding for a hypothetical protein with no putative conserved domains identified. These genes have the potential of affect the metabolism and the virulence of the bacterial strains. For example, although the major pilin synthesis gene is known to be characterised by locus variants which are associated with biofilm and virulence (namely variants PI-I, PI-IIa and PI-IIb, [Périchon *et al*., 2017]), CC17 is characterised by the presence of PI-I/PI-IIb. Smaller variations within the locus PI-IIb appear to be associated with CC17 isolated from carriage, suggesting that this gene may be impaired in functionality. Similarly, the *prtP* gene and the *glgD* genes, encoding respectively for a protease associated with virulence and for the ATP-binding cassette of a multidrug-efflux pump, have alleles that are more common in strains isolated from disease, highlighting the potential for these allelic variations to result in a more virulent phenotype [Obolski *et al*., 2019].

**Table 3 –.** CC17-associated genes showing at least one allele statistically associated with either strains isolated from invasive disease or from carriage. * = *gcc1730* shows only one mismatch in the protein alignment, which introduced a stop codon in position 122.

| Gene | Allele 1 |  | Allele 2 |  | Mismatches (aa) | % difference (aa) |
| --- | --- | --- | --- | --- | --- | --- |
|  | p-value | Odds-ratio | p-value | Odds-ratio |  |  |
| *gdh* | 1.16E-18 | 84.47 | 1.67E-03 | 0.19 | 0 | 0 |
| *dinG* | 1.95E-10 | 0.08 | 0.063 | 0.55 | 0 | 0 |
| *gcc178* | 3.83E-21 | 0.09 | 0.151 | 1.41 | 0 | 0 |
| *metN* | 6.07E-19 | 0.10 | 2.97E-14 | 0.10 | 1 | 0.4 |
| *yhjX* | 0.021 | 0.24 |  |  | 1 | 0.2 |
| *pta* | 0.013 | 0.19 |  |  | 1 | 0.3 |
| *strA* | 0.004 | 5.86 |  |  | 0 | 0 |
| *gcc1730* | 3.73E-03 | 5.93 |  |  | 1* | 0.3* |
| *gpp1725* | 6.43E-05 | 0.47 |  |  | 1 | 0.6 |
| *endA* | 4.07E-05 | 0.46 |  |  | 1 | 0.9 |
| *dtpT* | 1.25E-05 | 0.43 |  |  | 1 | 0.2 |
| *cadR* | 7.15E-06 | 0.42 |  |  | 1 | 0.3 |
| *pyrB* | 1.71E-08 | 0.10 |  |  | 1 | 0.3 |
| *gcc171* | 1.37E-09 | 0.15 |  |  | 1 | 1.1 |
| *gcc176* | 2.18E-10 | 3.56 |  |  | 1 | 0.2 |
| *gcc1713* | 1.87E-10 | 3.55 |  |  | 0 | 0 |
| *natA* | 3.02E-11 | 3.69 |  |  | 1 | 0.9 |
| *inlA_2* | 5.46E-16 | 0.09 |  |  | 1 | 0.1 |
| *prtP_2* | 3.31E-17 | 42.87 |  |  | 1 | 0.1 |
| *glgD* | 2.88E-18 | 83.77 |  |  | 1 | 0.9 |
| *efrB* | 1.15E-19 | 49.48 |  |  | 1 | 0.2 |

## DISCUSSION

*S. agalactiae* isolated from human and animal sources is characterised by a range of Clonal Complexes and Sequence Types. Each CC appears to be phenotypically different, with CC1 being commonly isolated in adult disease, and CC17 (associated with capsular serotype III) commonly isolated in neonatal disease and demonstrating hypervirulence [Teatero *et al*., 2017; Shabayek and Spellerberg., 2018]. We show that these different CCs are characterised by different gene sets belonging to functional families involved in niche adaptation and virulence. Furthmore, within CC17, we have identified several, functionally important alleic variants associated with either carriage or disease. We suggest that each human-associated CC has maintained these genes following zoonotic transfer [Botelho, *et al*., 2018]. This is in part reflected in the varying potential of different CCs to cause invasive diseases in different human hosts, as illustrated by the hypervirulence of CC17 in neonates, the lower neonatal invasive potential of CC1, CC19 and CC23 clones, and the propensity of CC1 to cause disease in adults with co-morbidities [Manning *et al*., 2008; Teatero *et al*., 2017]. Importantly, these CC-specific genetic characteristics and the pattern of gene presence and absence are independent of geographical origin, with the exception the CC10 gene cluster present in the strains isolated from Africa belonging to ST327 and 328.

Amongst the hypervirulent CC17-specific genes, there were several examples of previously identified genes associated with human disease due to GBS and other related bacteria. For instance, the transporter Nik which controls the uptake of nickel is essential for survival in the human host. A homologue of Nik has been shown to be essential for *Staphylococcus aureus* in the causation of UTIs [Remy *et al*., 2013]. The DLD gene, encoding for dihydrolipoamide dehydrogenase enzyme [Smith *et al*., 2002], has been implicated in several virulence related processes in *Streptococcus pneumoniae*, such as survival within the host and production of capsular polysaccharide. Mutants lacking the DLD gene are unable to cause sepsis and pneumonia in mouse models [Smith *et al*., 2002]. Surface proteases in *S. agalactiae* are described to have several virulence-associated functions, such as inactivation of chemokines that recruit immune cells at the site of infection or facilitate invasion of damaged tissue [Lindahl *et al*., 2005; Lalioui *et al*., 2005]. We have identified PrtP and ScpA proteases, both characterised by the presence of C5a peptidase domains and a signal peptidase SpsB, specific to this complex. Genes known to be associated with CC17 hypervirulence have also been identified in this analysis including the Pi-IIb locus [Périchon *et al*., 2017], part of which is represented by the CC17-associated genes *gcc1732, lepB, inlA_2, gcc1733* (Table S3), supporting the validity of this analysis. Allelic variation of virulence associated genes has previously been used to identify genes classifying invasive and non-invasive strains in other streptococcal species [Obolski *et al*., 2019]. A proportion of CC17 specific genes also showed unique alleles associated with invasive disease or carriage strains. Sixteen of 21 allelic variants resulted in a difference that was translated into the protein sequence, including regulatory proteins and virulence- or metabolism-associated proteins, such as ABC-transport systems, a major pilin protein and a C5a peptidase. These data suggest that there have been further selection processes within hypervirulent CC17 that could result in strains characterised by different virulence levels.

The CC23-specific genes identified are putatively involved in virulence and host invasion, including *mntH* a gene encoding for a manganese transport protein. During a bacterial infection the host limits access to manganese, amongst other micronutrients, and it has been shown that *S. aureus* responds to this host-induced starvation by expressing metal transporters, such as MntH [Kehl-Fie *et al*., 2013]. Interestingly, CC23 is also associated with *vanY*, a gene implicated in vancomycin resistance in other streptococci [Romero-Hernández *et al*., 2015]. GBS is typically susceptible to vancomycin [Berg *et al*., 2014], an antibacterial glycopeptide obtained from *Streptomyces orientalis* which inhibits cell wall synthesis, alters the permeability of the cell membrane and selectively inhibits ribonucleic acid synthesis [Moellering, 2005]. Whether the presence of this gene also facilitates niche adaptation in the context of complex host-microbiota environment remains to be determined.

Lineage CC10, and the sub-lineage CC19 that includes the strains belonging to the ST327 and ST328 are mostly characterised by metabolic genes, consistent with the lower virulence of these clonal complexes. The genes *galTKEM* are present in these two lineages only, and encode for the “Leloir pathway” in other streptococci, such as *mutans, thermophlus* and *pneumoniae* [Vaillancourt *et al*., 2002; Abranches *et al*., 2004; Anbukkarasi *et al*., 2014]. This pathway in *S. pneumoniae* is finely tuned by CbpA, and activated in tandem with the tagatose-6-phosphate pathway in order to maximise growth [Carvalho *et al*., 2011]. The functionality of this pathway is yet to be described in GBS, but we hypothesise that accessing different methods to metabolise carbohydrates facilitates nutrient competition and survival. Amongst the non-metabolic genes that are associated with CC19, we identified a *virD4* gene, which is part of a previously identified type IV secretion system (T4SS) [Zhang *et al*., 2012] present in numerous bacterial species and associated with virulence effector translocation and conjugation [Alvarez-Martinez and Christie, 2009; Wallden *et al*., 2010].

GBS is widely thought to be a zoonosis [Botelho *et al*., 2018; Zadoks *et al*., 2011; Lyhs *et al*., 2016; Manning *et al*., 2010; Chen *et al*., 2015]. Based on the CC-characterising genes that we identified, their relative frequency in the GBS population, and their distribution in the GBS genome, we hypothesise that *S. agalactiae* lineages that colonise humans initially evolved in animals and then subsequently expanded clonally in humans. In line with the observation that *S. agalactiae* has undergone genome reduction [Rosinski-Chupin *et al*., 2013], we suggest that the human-adapted clones evolved in animals through loss of function of redundant genes. Having escaped the animal niche, they were then able evade the human immune system and establish successful colonisation (Figure 4). Recently, the “missing link” between animal and human adaptation of GBS was described to be CC103 [Botelho *et al*., 2018]. However, we have identified animal isolates belonging to human-associated CCs (e.g. CC17 and CC23) which cluster together with human clinical isolates in the GBS population structure.

**Figure 4 –.**
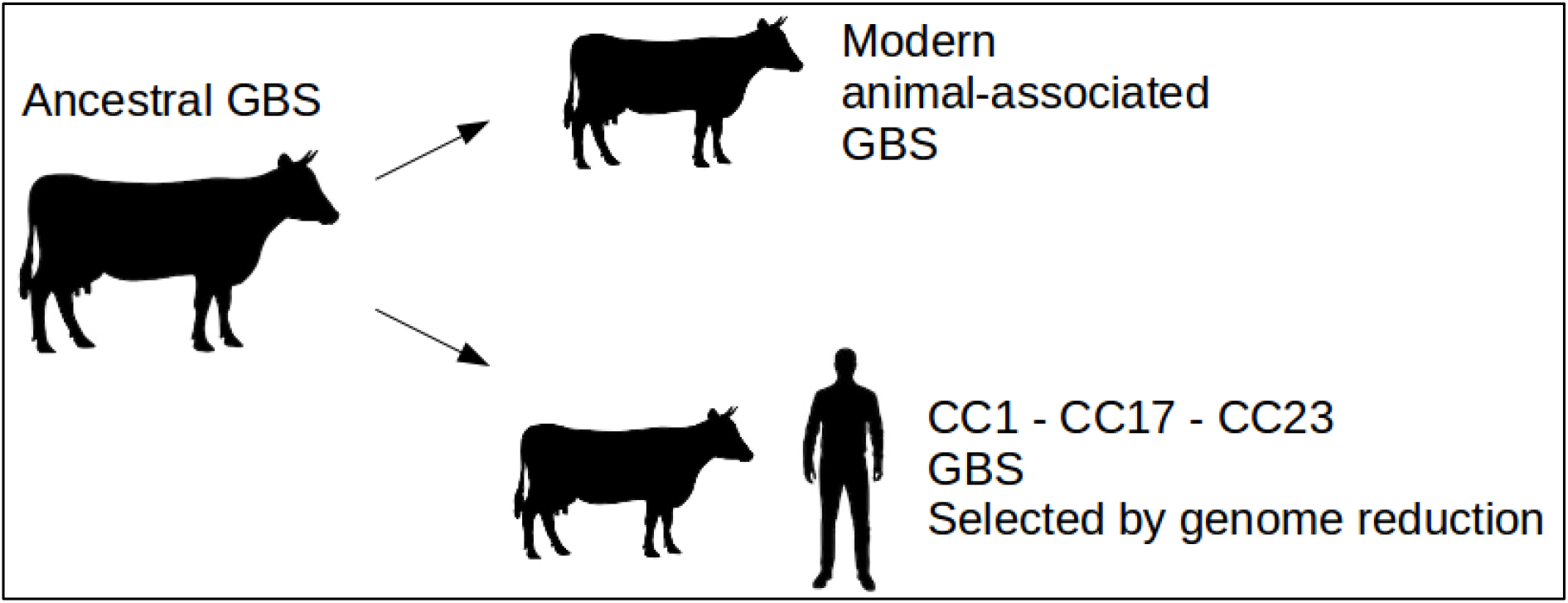
Hypothesis: a model for the differentiation of GBS into animal- and human-associated strains. The ancestral GBS strains carried every gene present in each of the modern clonal complex. Modern animal strains, cluster in a single clade characterised by a very high variability and deep branching. Modern human and animal strains belonging to CC1 (including CC10 and CC19), CC23 and CC17 have differentiated by genome reduction and clonal expansion. Hypervirulent clones (e.g. CC17) retained genes useful for the colonisation of the human niche.

Our analysis has a number of limitations. Firstly, we were confined to the current publicly available GBS human and animal genomes retrieved from pubmlst.org/sagalactiae/ (a total of 3028 isolates including the full dataset from The Netherlands), plus a further 303 genomes from Malawi. Secondly, the GWAS pipeline we used relies on the automated annotation of software Prokka. The use of this software required the use of Roary and Scoary to produce the pangenome and the pan-GWAS. This was extremely efficient when used to annotate the thousands of bacterial genomes in this analysis, and although the genome annotations and the pan-genome were manually screened for consistency and quality (such as saturation of the core and accessory genome), it could potentially introduce artefacts. Confirming the GWAS findings with the sequence alignments allowed us to identify several genes that were characterised by non-synonymous mutations and small in-dels, as well as it unravelled these potential artefacts that require further investigation. Finally, our analysis is confined to the genomic differences between the different clades, further laboratory and epidemiological analysis will be needed to fully appreciate the biological consequences of these CC-specific genes.

In conclusion, we have shown that the CCs of *Streptococcus agalactiae* responsible for neonatal meningitis and adult colonisation are characterised by the presence of specific gene sets that are not limited to particular geographical areas. We suggest that human-associated GBS CCs have largely evolved in the animal host before spreading clonally to the human, enabled by functionally different sets of CC-specific genes which enable niche adaptation. In the context of GBS control measures such as vaccination, we speculate that as the human gastrointestinal and urogenital niches are vacated by vaccine serotypes, serotype-replacement could occur as a result of new GBS strains arising from animals including cattle and fish, reservoirs of GBS genetic diversity.

## ACKNOWLEDGEMENTS

The authors would like to thank all the clinical and laboratory staff at the MLW Clinical Research Programme in Malawi and all families and their infants who participated in the study.

## FUNDING

This work was funded by a project grant from the Meningitis Research Foundation (Grant 0801.0); a project grant jointly funded by the UK Medical Research Council (MRC) and the UK Department for International Development (DFID) under the MRC/DFID Concordat agreement and is also part of the EDCTP2 programme supported by the European Union (MR/N023129/1); and a recruitment award from the Wellcome Trust (Grant 106846/Z/15/Z).

The MLW Clinical Research Programme is supported by a Strategic Award from the Wellcome Trust, UK. The NIHR Global Health Research Unit on Mucosal Pathogens at UCL was commissioned by the National Institute for Health Research using Official Development Assistance (ODA) funding.

## DISCLAIMER

The views expressed are those of the author and not necessarily those of the NHS, the NIHR or the Department of Health and Social Care.

## SUPPLEMENTARY MATERIAL

**Table S1 – Metadata of each GBS isolate described in this work.** From left to right each column shows the isolate name, the clonal complex to which the isolate belongs (CC1, CC6, CC10, CC19, CC17, CC23, Single ST or N/D), the source of isolation (animal or human, in which case it is reported as carrier, invasive or unknown), the country of isolation, the serotype, the year of isolation (where available), the MLST-type, and the accession number of each isolate (where available).

**Table S2 – Sequence type defining each clonal complex and number of STs isolated per country.** The table is divided in 8 horizontal sectors (CC1, CC6, CC10, CC19, CC17, CC23, Single ST or N/D). In each sector columns show (from left to right) which ST is represented in each CC, number of isolates belonging to a particular ST are found in each of the country where the isolated described in this study were sourced (Brazil, Canada, Germany, Italy, Kenya, Malawi, Netherlands, USA or unknown source).

**Table S3 – Genes defining each CC as identified by pan-GWAS.** The table is divided in five sections, relative to CC1, CC10, CC19, CC17 and CC23. Each column shows (from left to right) the name of the gene identified by pan-GWAS, the length of the putative protein produced by each gene, the KEGG database id (where available), a short annotation of each gene (according to KEGG and/or Prokka where available, otherwise reported as hypothetical protein), the functional class to which each gene belongs (metabolic – reported as “met”, environmental information processing - “env” or cellular processes - “cell”, according to KEGG annotation, the type of variation between different clonal complexes (“Point mutations”, “Synonymous point mutations”, “Truncated protein”, “Gene absent”).

In case both prokka and KEGG annotation did not report a gene name, an arbitrary gene name was assigned to the hypothetical gene following the scheme, “g”, followed by the clonal complex and an incremental number.

**Figure S1 - Core genome based population structure of GBS built with alternative reference strains.** Trees showing the GBS population structure as in figure 1, produced with a different reference strain. Reference strains used for the four trees are: NC_021485 – strain 09mas018883; NC_007432 – strain A909; NC_018646 - strain GD201008; NC_004368 – strain NEM316. For each tree, the annotation is analogous to the one described in figure 1.

**Figure S2 – Heatmaps based on the BLASTn score of each CC-characterising gene for each isolate.** Heatmaps were produced with pheatmap package in R (clustering of rows and column was performed using Euclidean method). Each heatmap shows 1988 isolates on the rows and the CC-associated genes on the column. Each row-clustering tree (related to the isolates) is annotated with coloured strips representing the Clonal complex. Strains belonging to CC10-ST327 and CC10-ST328 are reported as CC327 in this representation

**Figure S3 – Syntheny of CC17 characterising genes.** The image shows the alignment of three CC17 *S. agalactiae* genomes (strain 874391, BM110 and SGM6) and 104 CC17-associated genes (at the bottom). Each vertical line represents a sequence that is found in the same location in all the analysed sequence. Image obtained with software Mauve.

**Figure S4 –Genes showing alleles statistically associated with carriage or invasive disease inCC17 strains.** Each barplot shows the frequency of each allele in each of the 21 CC17-associated gene observed to have at least one allele associated with disease or carriage. Different numbers on the x-axis represent different allelic configuration of the gene. P-value < 0.05. * = Non-significant.

**Supplementary File 1 – Representative sequences CC-specific genes.**

## Notes

#### Summary of Updates

Section "Functional pathways affected by CC-specific genes" updated with a paragraph on CC17-specific alleles. Table 3 and Figure S4 added.

