## Supplementary figures and images for "Pan-GWAS of *Streptococcus agalactiae* highlights lineage-specific genes associated with virulence and niche adaptation"

### Supplemental Figure 1

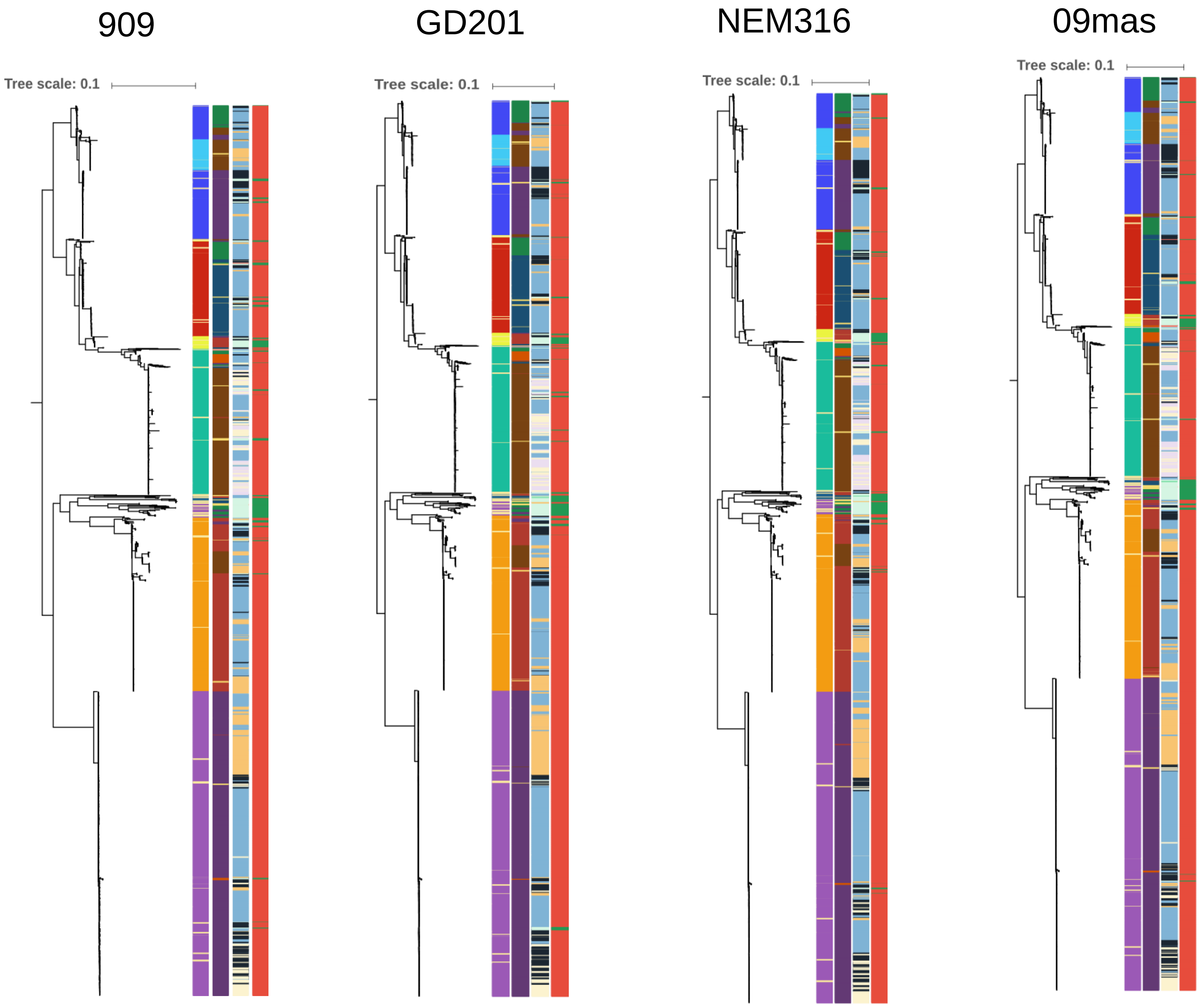

### Supplemental Figure 2

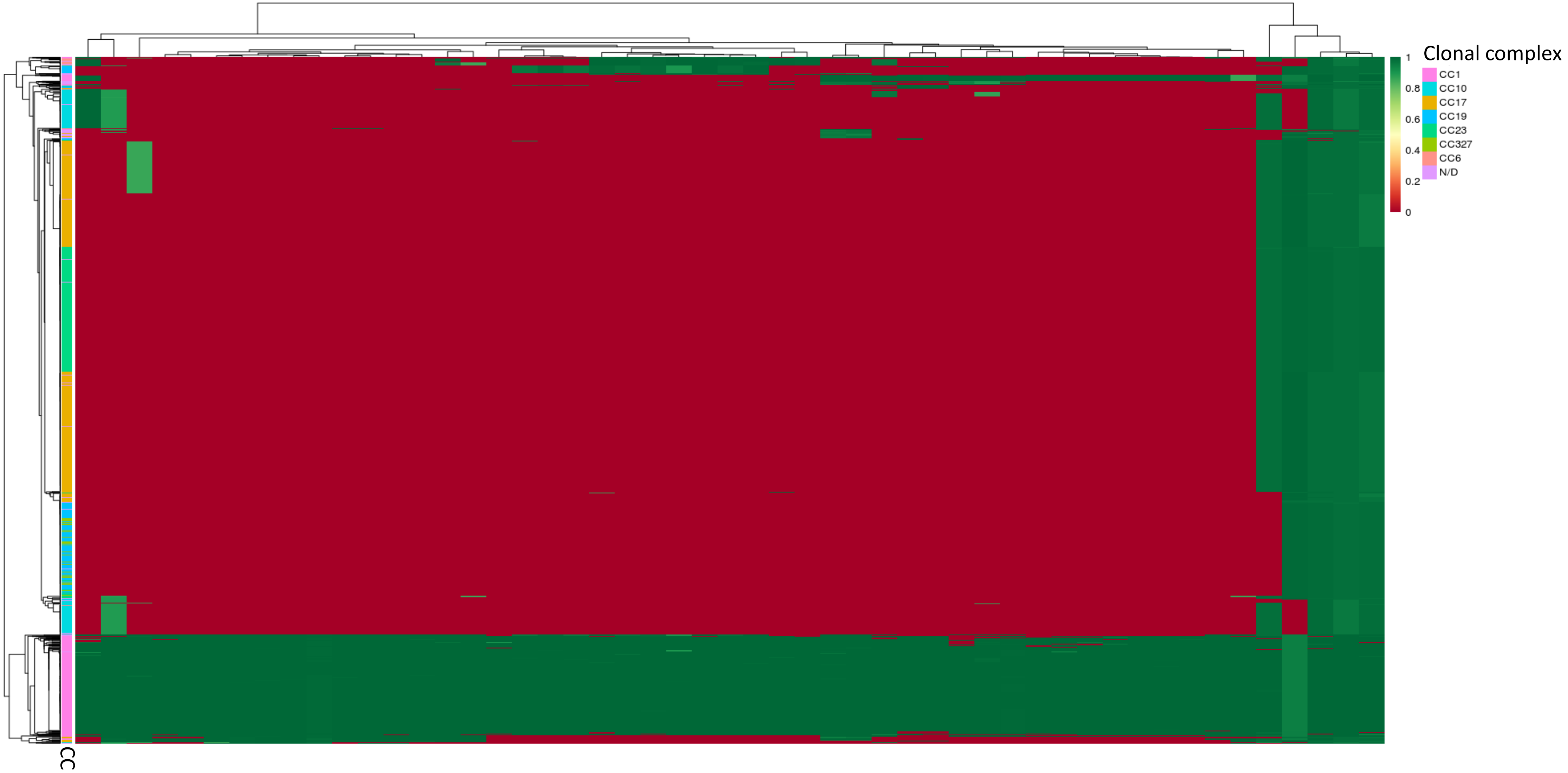

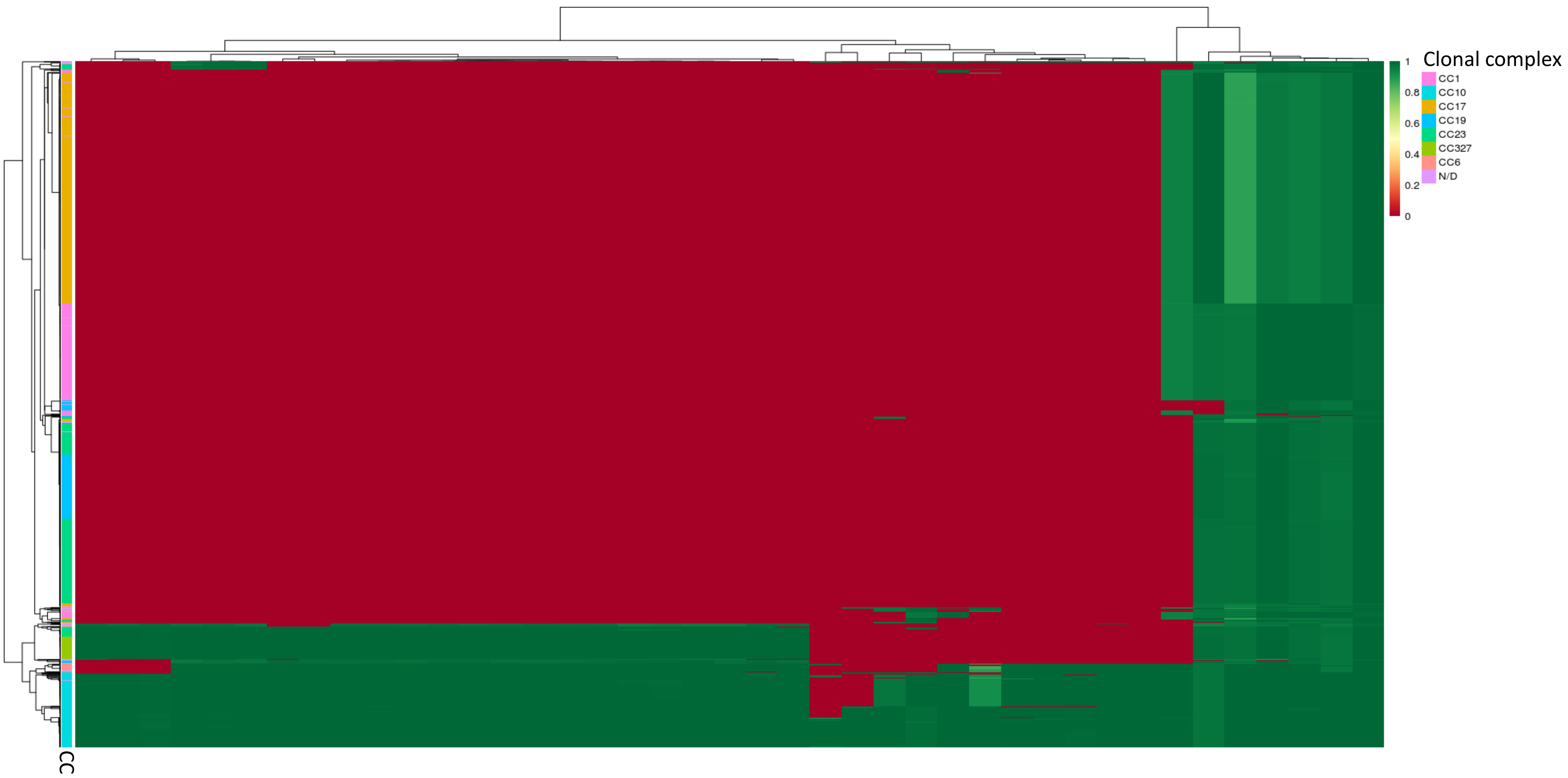

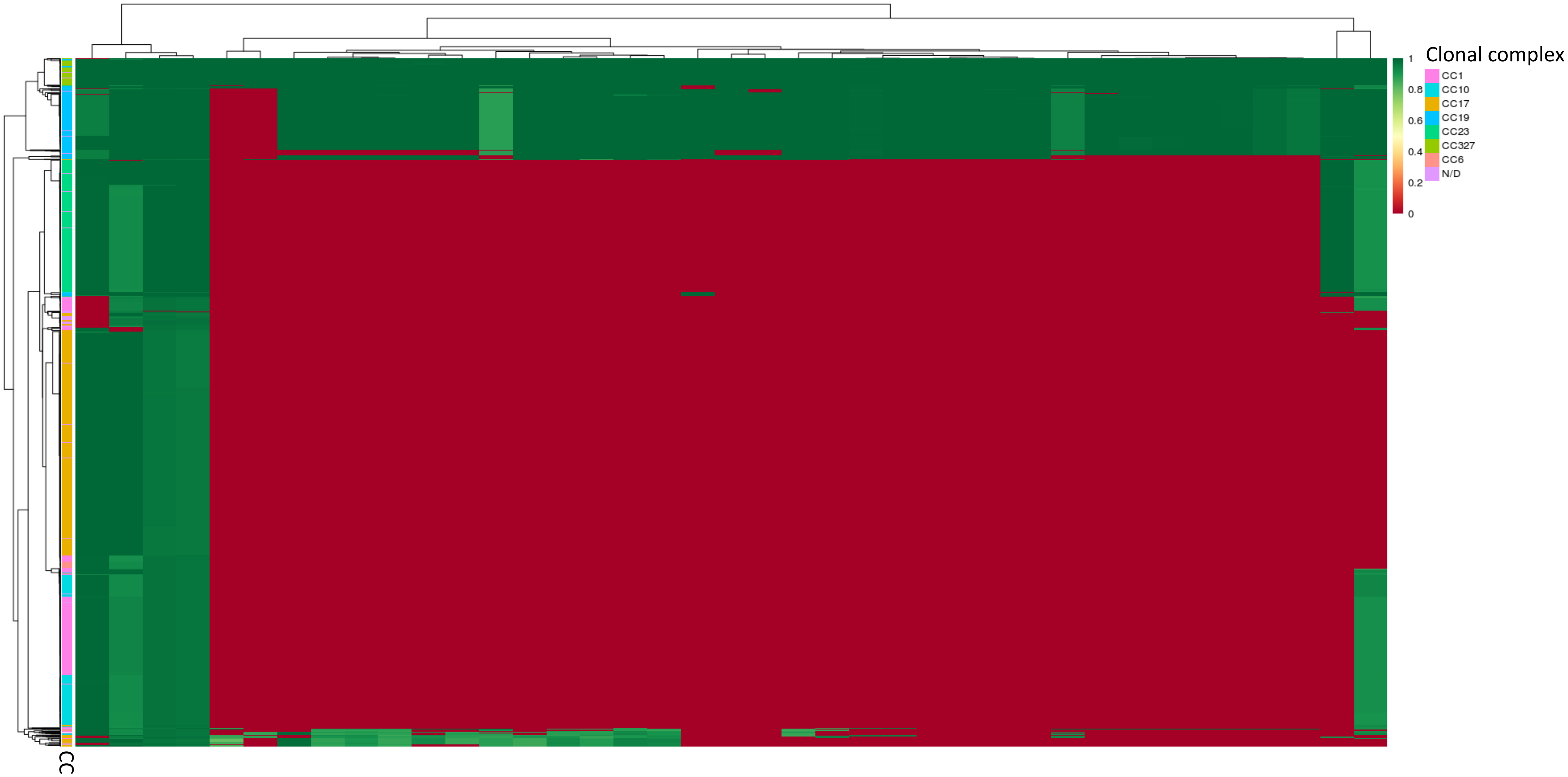

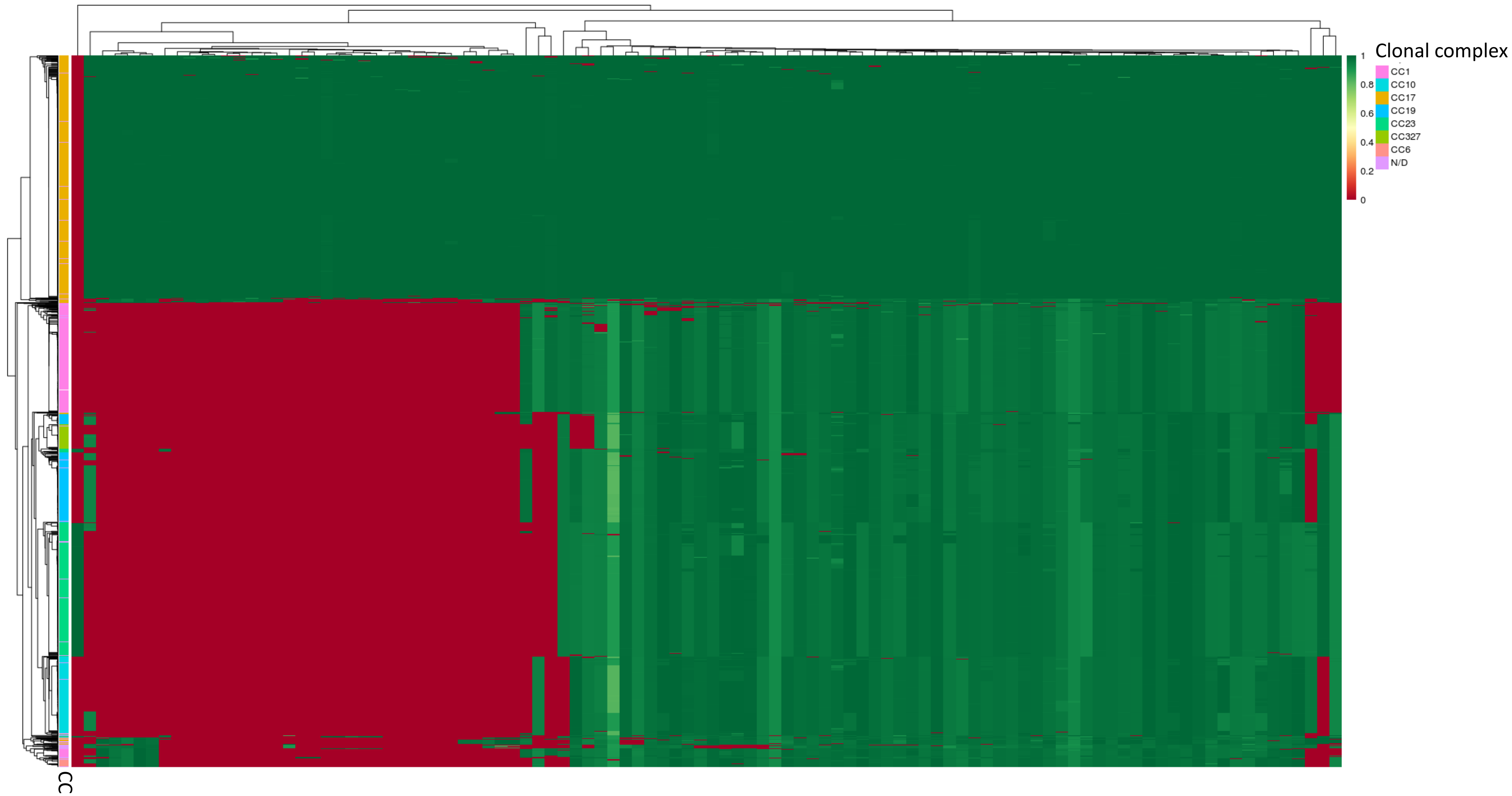

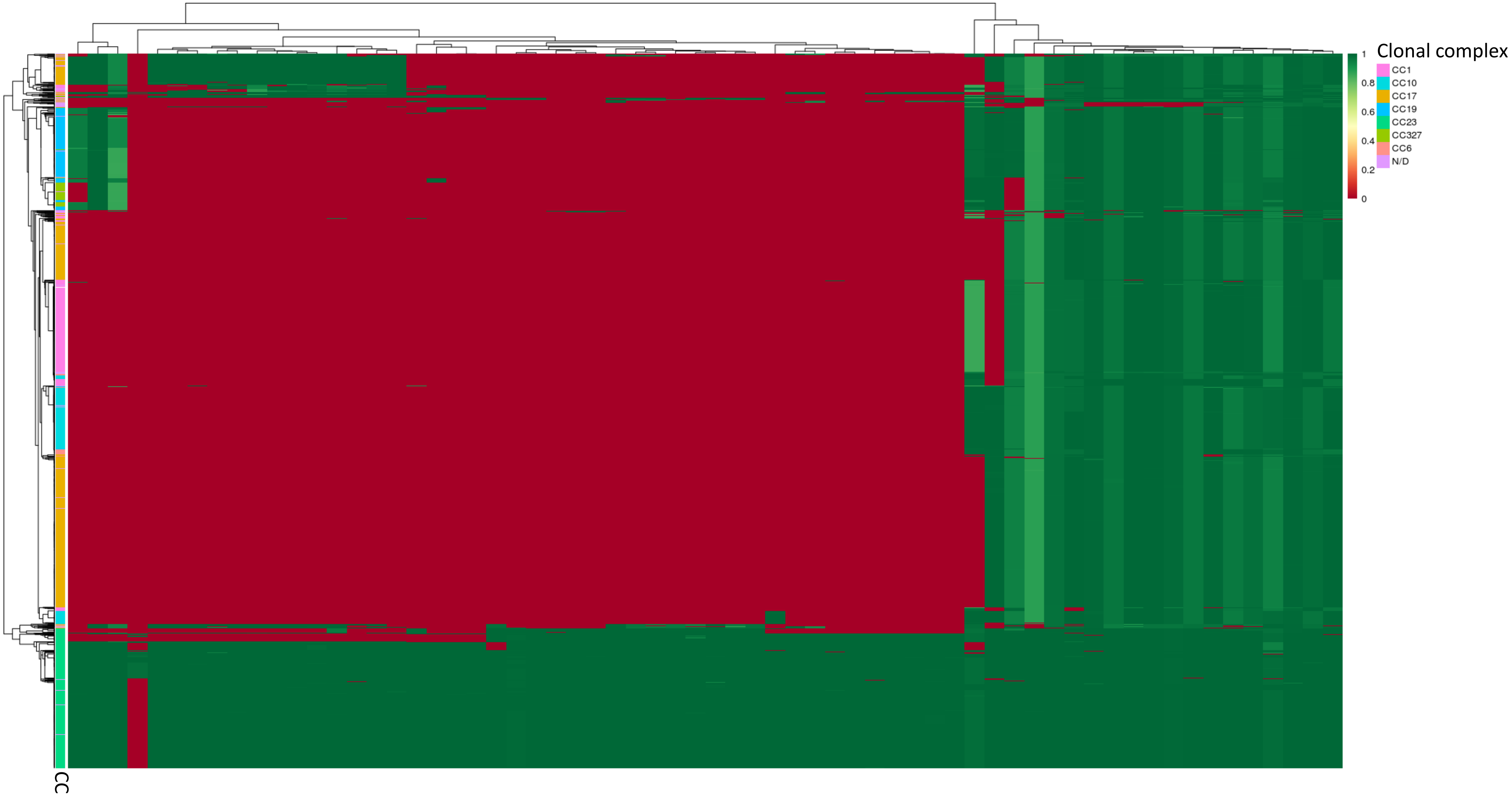

### Supplemental Figure 3

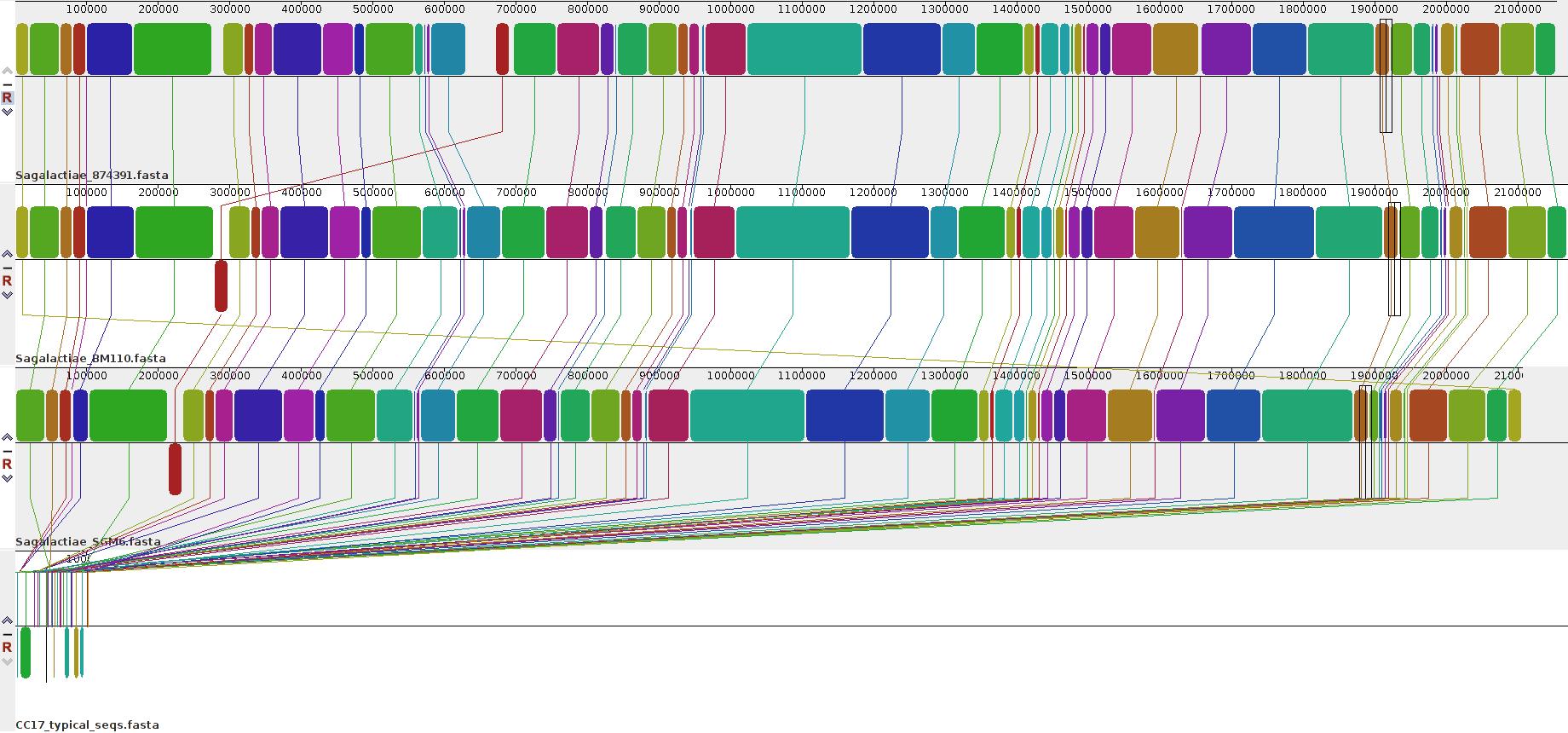

### Supplemental Figure 4

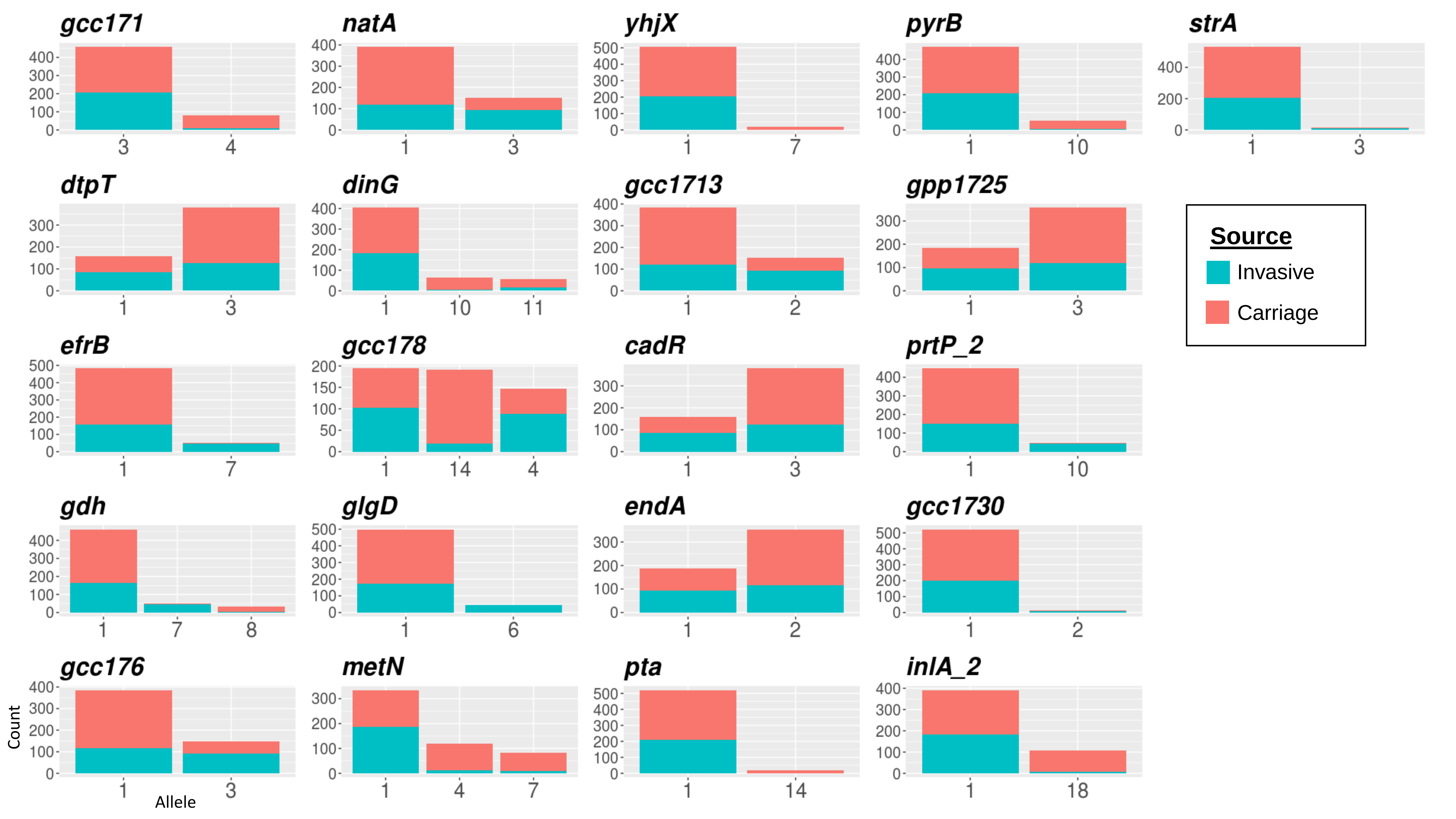
